# Balance between extracellular matrix production and macrophage survival by a *Salmonella*-specific SPI-2 encoded transcription factor

**DOI:** 10.1101/2021.03.09.434593

**Authors:** María Laura Echarren, Nicolás R. Figueroa, Luisina Vitor-Horen, M. Graciela Pucciarelli, Francisco García-del Portillo, Fernando C. Soncini

## Abstract

Cellulose is a major component of the *Salmonella* biofilm extracellular matrix and it is considered an antivirulence factor because it interferes with *Salmonella* survival inside macrophages and virulence in mice. Its synthesis is stimulated by CsgD, the master regulator of biofilm extracellular matrix formation in enterobacteria, which in turn is under the control of MlrA, a MerR-like transcription factor. In this work we identified a SPI-2 encoded *Salmonella*-specific transcription factor homolog to MlrA, MlrB, that represses transcription of its downstream gene, STM1389, also known as *orf319*, and of *csgD* inside host cells. MlrB is induced in laboratory media mimicking intracellular conditions and inside macrophages, and it is required for intramacrophage survival. An increased expression of *csgD* is observed in the absence of MlrB inside host cells. Interestingly, inactivation of the CsgD-controlled cellulose synthase coding-gene, *bcsA*, restored intramacrophage survival to rates comparable to wild type bacteria in the absence of MlrB. These data indicate that MlrB represses CsgD expression inside host cells and in consequence activation of the cellulose synthase. Our findings provide a novel link between biofilm formation and *Salmonella* virulence.

## INTRODUCTION

*Salmonella enterica* success on infection and transmission between hosts relies greatly on its ability to recognize, to evade and to even exploit host defenses in its favor (Behnsen *et al*., 2015). Critical to *Salmonella* pathogenesis is the coordinated assembly of two type III secretion systems (T3SS) which are located at specific genome regions called pathogenicity islands (SPI). The *Salmonella* pathogenicity island 1, SPI-1, is associated with the invasion of epithelial cells, whereas SPI-2 is required for intracellular survival (Deiwick *et al*., 1999; Ellermeier and Slauch, 2007; McGhie *et al*., 2009; Agbor and McCormick, 2011), although there is certain overlap in their functions (Deiwick *et al*., 1998; Steele-Mortimer *et al*., 2002; Brown *et al*., 2005). Aside from these orthodox virulence traits, it is widely recognized that *Salmonella* resistance in extra-host environments is also an important factor for the attainment of the infectious cycle (Waldner *et al*., 2012; Maruzani *et al*., 2019). Essential to the interaction with hosts and the environment is its ability to form biofilms, bacterial communities embedded in an adhesive, self-produced extracellular matrix (EM) with an intricate physiological and structural organization (Serra *et al*., 2013).

Although accumulated evidence supports the importance of biofilms in environmental conditions, the role of the community lifestyle in *Salmonella* pathogenic processes is far less understood (Maruzani *et al*., 2019). Most host-generalists involving non-typhoidal serovars preserve their ability to form biofilms *in vitro* (reviewed in MacKenzie *et al*., 2017). For these pathovars, it has been suggested that the biofilm formation not only favors resistance inside the gastrointestinal tract (MacKenzie *et al*., 2017), but it also modulates the local inflammatory responses, contributing to the pathogen competitiveness and, therefore, transmission (Thiennimitr *et al*., 2012). Host-specific typhoidal serovars, on the other hand, are able to adapt the expression and composition of their EM avoiding host immunity (Gonzalez-Escobedo and Gunn, 2013; Adcox *et al*., 2016). In a broader context, it is increasingly accepted that biofilms promote long-term, intra-host persistence in a variety of pathogenic bacteria, whereas planktonic lifestyle favors acute infection (Crawford *et al*., 2010; Desai and Kenney, 2019). In line with the latter, overproduction of cellulose, one of the major *Salmonella* EM components, has been proven to hinder virulence (Pontes *et al*., 2015; Ahmad *et al*., 2016). Nevertheless, it has also been found that basal expression levels of cellulose are present when inside macrophages (Pontes *et al*., 2015). These findings support a role of EM production in the regulation of virulence, although much remains to be explored about the molecular mechanisms relating *Salmonella* community lifestyle and the infectious process.

In *Salmonella* and related enterobacteria, transition to the multicellular behavior is primarily controlled by the master regulator CsgD that activates the synthesis of the major EM components, curli and cellulose (Gerstel and Römling, 2003). CsgD expression is modulated by several factors that integrate diverse environmental signals into what is today recognized as one of the most complex bacterial regulation networks known (Gerstel and Römling, 2003; Ogasawara *et al*., 2011). Among those factors, MlrA, a MerR-like family response regulator, has been identified as a key *csgD* transcriptional activator in *Escherichia coli* and *S.* Typhimurium (Brown *et al*., 2001). MlrA recognizes a symmetrical dyad sequence, its operator, ∼100 bp upstream of the -35 element in the *E. coli csgD* promoter (Ogasawara *et al*., 2010), and a similar operator sequence is distinguished in the *Salmonella csgD–csgB* intergenic region.

In this work, we detected the presence of a *Salmonella*-specific, MerR-like transcription factor, Orf242, which we renamed MlrB for *<u>M</u>erR-like <u>r</u>egulator <u>B</u>*, that shows sequence homology to MlrA, specially at the DNA recognition domain. We identified that MlrB acts as a repressor of both *csgD* and the *mlrB* downstream gene, STM1389, also known as *orf319*. MlrB is not only induced in media mimicking intracellular conditions, but it is also required for *Salmonella* intramacrophage survival. An increased expression of *csgD* is observed in the *mlrB* deleted strain indicating that it exerts this function by reducing *csgD* expression inside host cells. These results provide a novel link between biofilm formation and *Salmonella* virulence. Additionally, they allow us to postulate a mechanism by which two close homologues act antagonistically depending on the specific niche.

## RESULTS

### Identification and genetic context of MlrB, a *Salmonella*-specific MlrA-like transcriptional regulator

*STM1390*, also known as *orf242*, encodes a putative, *Salmonella-*specific MerR-like family protein of 242 amino acid residues, similar to MlrA, a key regulator controlling *csgD* expression and production of extracellular matrix (Brown *et al*., 2001) Conservation between these two regulators (40% amino acid sequence identity, and 70% similarity) becomes more evident at their N-terminal domains (Fig. S1A), especially on the residues purportedly involved in DNA recognition (Brown *et al*., 2001). Sequence divergence between MlrA and MlrB is higher at their C-terminals, which is a region typically involved in signal recognition among members of the MerR family of transcriptional regulators (Brown *et al*., 2003). In particular, as demonstrated for *E. coli*, the C-terminal domain of MlrA exerts a regulatory role by interacting with proteins of the c-di-GMP signaling pathway (Lindenberg *et al*., 2013). Regarding *STM1390* (hereinafter “*mlrB”*), inference of its physiological function through homology search remains elusive, as this region shows no similarity to any known protein domain.

The *mlrB* coding gene is located within the *Salmonella* SPI-2 (Fig. S1B), which is a species-specific, chimeric genomic island with two distinctive DNA fragments acquired by different horizontal transfer events (Hensel, Nikolaus, *et al*., 1999; Hensel, 2000). One encodes the majority of structural, regulatory and effector factors of the Type III Secretion System (T3SS-2) involved in *Salmonella* intracellular survival during infection (Hensel, 2000). The other encodes the tetrathionate reductase system involved in the outgrowth of *Salmonella* during the intestinal colonization (Hensel, Hinsley, *et al*., 1999; Winter *et al*., 2010) as well as various genes of yet unknown function, including *mlrB*.

Considering the similarity between MlrB and MlrA, and *mlrB* location in the genome, we investigated whether MlrB also modulates EM production, and whether it has a role in *S.* Typhimurium virulence.

#### MlrA, rather than MlrB promotes biofilm formation and *csgD* transcription under low osmolarity

We first analyzed a potential role of MlrB in EM production by evaluating biomass adhesion to an abiotic surface using crystal violet-binding quantification (see Materials and Methods, and Pitts *et al*., 2003). Biomass adherence of strains deleted in either *mlrA*, *mlrB* or the *mlrA mlrB* double mutant was compared after incubation at 28°C for 48 h in LB broth lacking sodium chloride (SLB). These conditions were reported to promote *in vitro* biofilm formation in *Salmonella* via activation of *csgD* expression (Kader *et al*., 2006). Regardless the sequence similarity between these two regulators, the Δ*mlrB* strain showed no difference from the wild-type strain either in the amount of adhered biomass (Fig. 1A) or in *csgD* expression (Fig. 1B). A possible explanation for this result is that MlrB deficiency in the Δ*mlrB* strain could be masked by the presence of MlrA. However, the low and similar *csgD* expression observed in both Δ*mlrA* and Δ*mlrA* Δ*mlrB* mutants (Fig 1B), along with the lack of adhesion of the Δ*mlrA* strain (Fig 1A) suggest otherwise. More likely, the conditions in which the experiments were carried out might not be optimal for MlrB activity, and thus a dominant role of MlrA is observed. In sum, under low osmolarity, MlrA exerts a predominant role in controlling of biofilm formation and *csgD* expression over its homologue MlrB.

**Figure 1.**
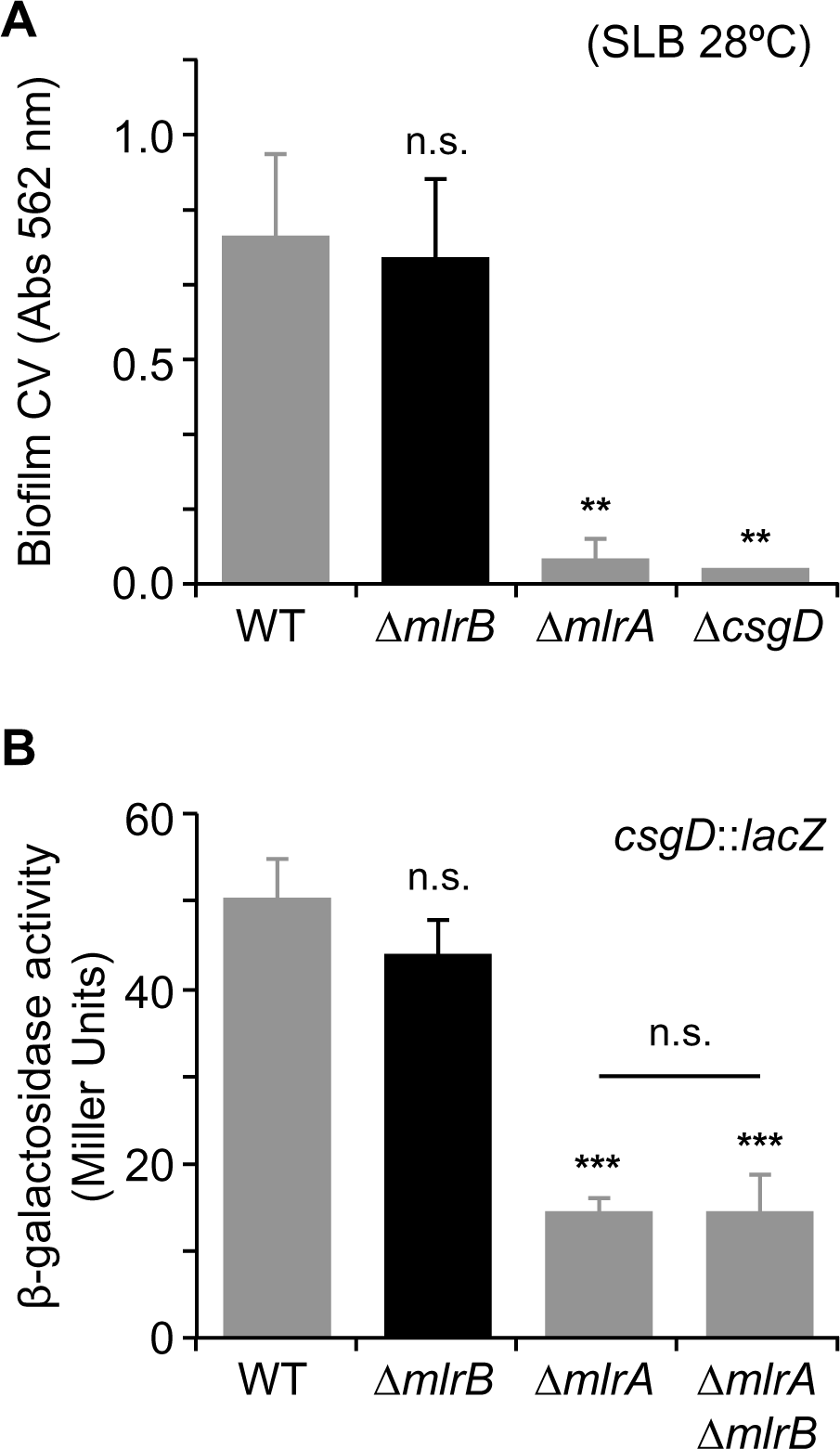
MlrB is not affecting CsgD expression or biofilm formation under laboratory conditions. **A.** Adherence to polystyrene. 1:100 dilutions of ON SLB cultures of the indicated strains were incubated in SLB medium, at 28 °C for 48 hours in polystyrene microplates without shaking. After successive washes, staining was performed with 1% crystal violet and 562 nm Absorbance was determined. The data correspond to average values of three independent experiments carried out by quadruplicate. The error bars correspond to the SD. **B.** β-galactosidase activity from a *csgD*::*lacZ* transcriptional fusion determined for wild-type (WT), Δ*mlrB*, Δ*mlrA* or Δ*mlrA* Δ*mlrB* cells, grown overnight in LB medium, at 28 °C. The data correspond to mean values of three independent experiments performed in duplicate. Error bars represent SDs. Symbols above bars denote statistical differences between means, with respect to WT. n.s., not significant; *, P < 0.05; **, P < 0.01; ***, P < 0.001.

#### *mlrB* is induced in SPI-2 *in vitro* conditions and inside macrophages

*mlrB* is located immediately adjacent to and downstream of *ssrB* (Fig. S1B), the gene coding for the response regulator of the SsrAB two-component system that controls the transcription of the *Salmonella* T3SS-2 (Deiwick *et al*., 1999; Walthers *et al*., 2007). We examined whether *mlrB* expression is coordinated with the rest of the T3SS-2 genes and whether this depends on SsrB. *mlrB* transcription increases under SPI-2-inducing conditions, i.e., PCN, InSPI-2 (Kröger *et al*., 2013) or LPM (Coombes *et al*., 2004), compared to LB (Fig. 2A and Fig. S2A) whereas *mlrA* transcription showed an opposite pattern (Fig. 2A and Fig. S2B). Using the strains harboring the *mlrB*::3xFLAG and *mlrA*::3xFLAG alleles tagged at their 3’ end, we determined that MlrB levels also had a 2.4-fold increment in InSPI-2 medium while MlrA had a 3-fold decrease, compared to LB (Fig. 2B). Although our results showed that the expression of MlrB follows that of T3SS-2 genes, deletion of *ssrB* had no effect on *mlrB* transcription (Fig. S3A) indicating that *mlrB* is neither co-transcribed with *ssrB*, nor is under SsrB transcriptional control. Conversely, deletion of MlrB had no effect on the transcription of *sseA* (Fig. S3B), a SsrA/SsrB regulated gene (Hensel, Nikolaus, *et al*., 1999; Hensel, 2000), suggesting that MlrB does not influence the SsrA/ssrB regulon.

**Figure 2.**
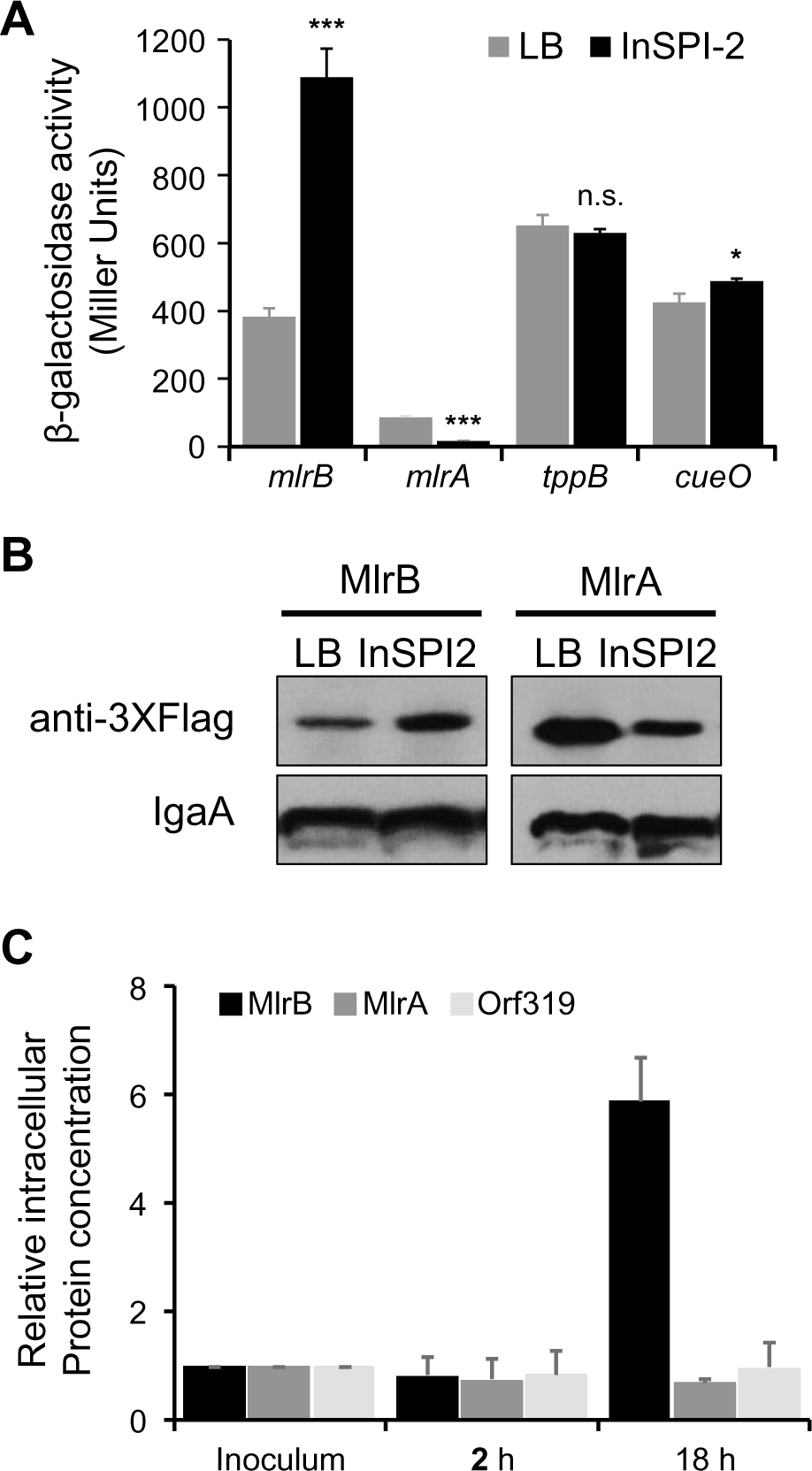
MlrB expression increases inside macrophages. A.β-galactosidase activity determined for the indicated reporter strains, grown until stationary phase in LB medium (gray bars) or InSPI-2 medium (black bars); values reported are means and SD of three independent experiments. Symbols above bars denote statistical difference between means from LB *versus* InSPI-2-grown cells. n.s., not significant; *, P < 0.05; ***, P < 0.01; ***, P < 0.001. **B.** Western blot analysis of MlrB and MlrA proteins labeled with the 3xFLAG epitope. Whole cell extracts were prepared from stationary phase cultures grown in the indicated media. Prior to processing, OD_600_ nm of each sample was adjusted to 1. Detection of IgaA with Anti- IgaA polyclonal antibodies was used as load control. **C.** Relative concentration of MlrB and MlrA determined from *Salmonella* cells grown inside macrophages at the indicated times. Concentrations were estimated from digitized autoradiographs by pixel densitometry. Original WB obtained from two independent experiments are included in Fig. S4.

We then determined the relative abundance of the 3xFLAG-tagged MlrB and MlrA from *Salmonella* cells grown inside RAW 264.7 macrophages, using anti IgaA as control. In accordance to the previous results we observed that MlrB increased almost 6-fold at 18 h.p.i. while we did not detect a substantial modification in MlrA concentration (Fig. 2C and S4).

#### *orf319* is a SPI-2 encoded- MlrB-repressed gene

We searched for putative MlrB-regulated genes in the *Salmonella* genome taking advantage of the similarity between MlrB and MlrA DNA-binding domains (Fig. S1A). Using the reported MlrA-recognized sequences in *E. coli* (Ogasawara *et al*., 2010), a positional weight matrix was generated with the MEME tool (Bailey *et al*., 2009). The resulting motif was confronted to the *S.* Typhimurium LT2 genome using the FIMO tool (Grant *et al*., 2011), also part of MEME suite. A sequence similar to the MlrA-consensus operator was identified by this method in the *csgD-csgB* intergenic region, 149 bp upstream of the predicted *csgD* transcription start site (Fig. S5). In addition, we detected a second putative MlrA operator, located 113 bp upstream of the *csgD* transcription start site (Fig. S5). Additionally, an MlrA-like operator was detected in *orf319* promoter region, a SPI-2 gene coding for a protein of unknown function, located immediately downstream of *mlrB* (Fig. S1B). This sequence was located at only 6 bp upstream of the predicted *orf319* transcriptional start site, overlapping the -10 and -35 promoter elements (Fig. S5B).

We tested whether *orf319* is under the transcriptional control of either MlrA or MlrB. Although both regulators repressed *orf319* transcription when *Salmonella* was grown in LB (Fig. S6), and their simultaneous deletion further increased the transcription of the target gene, only MlrB affected *orf319* transcription under SPI-2 inducing conditions (InSPI-2, Fig. 3). Furthermore, in InSPI-2 conditions, the Δ*mlrB* Δ*mlrA* double mutant showed an *orf319* transcription level similar to that displayed by the Δ*mlrB* strain. This correlates well with the reduced expression of MlrA as well as the increased expression of MlrB in this condition (Fig. 2 and S2).

**Figure 3.**
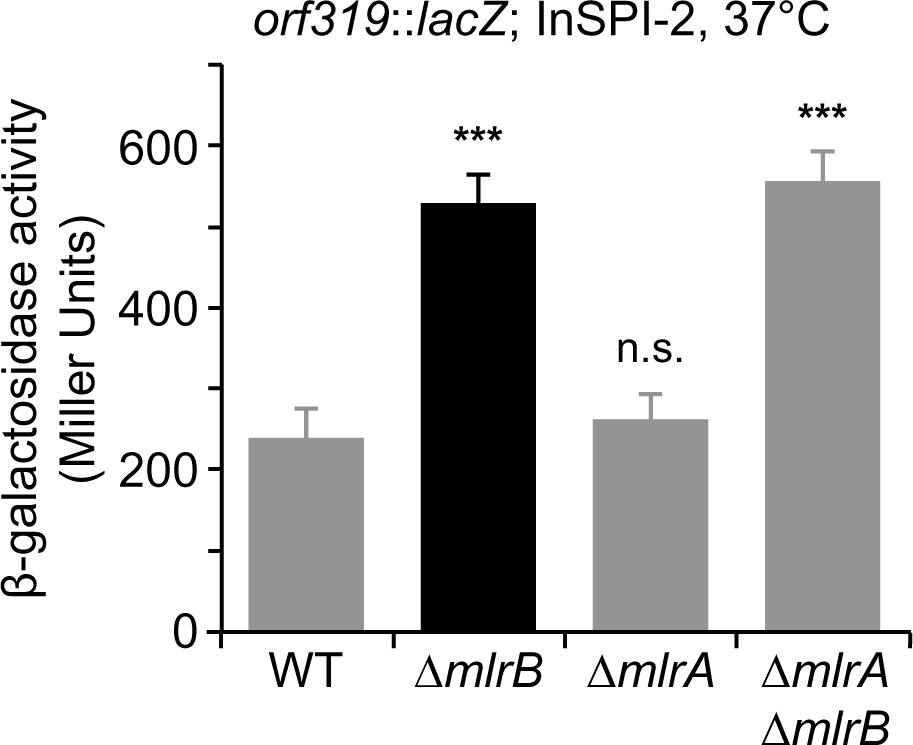
MlrB regulates *orf319* transcription under SPI-2-inducing conditions. β-galactosidase activity from an *orf319*::*lacZ* transcriptional fusion expressed on wild-type (WT), Δ*mlrB*, Δ*mlrA* and Δ*mlrA* Δ*mlrB* cells grown overnight in InSPI-2 medium at 37°C. The data correspond to mean values of tree independent experiments performed out in duplicates. Error bars represent SD. Symbols above bars denote statistical differences between means, with respect to WT. n.s., not significant; *, P < 0.05; **, P < 0.01; ***, P < 0.001.

#### MlrB is required for intramacrophage survival

MlrB increased expression inside macrophages and under SPI-2 inducing conditions, prompted us to study whether this regulator is required for intramacrophage survival.

Similar to mutants in the T3SS-2 system (Cirillo *et al*., 1998), the Δ*mlrB* strain had a defect in replication inside RAW 264.7 macrophages compared to the WT (Fig. 4A). Interestingly, MlrB overexpression from a multi-copy plasmid increased survival compared to the WT strain (Fig. 4B). Conversely, the Δ*mlrA* strain showed higher survival than the wild-type strain (Fig 4A), and its overexpression decreased *Salmonella* intramacrophage survival (Fig. 4B). Consistent with the opposite effect of both regulators on intracellular survival, the Δ*mlrA* Δ*mlrB* double mutant strain showed a survival behavior closed to the wild-type (Fig. 4A).

**Figure 4.**
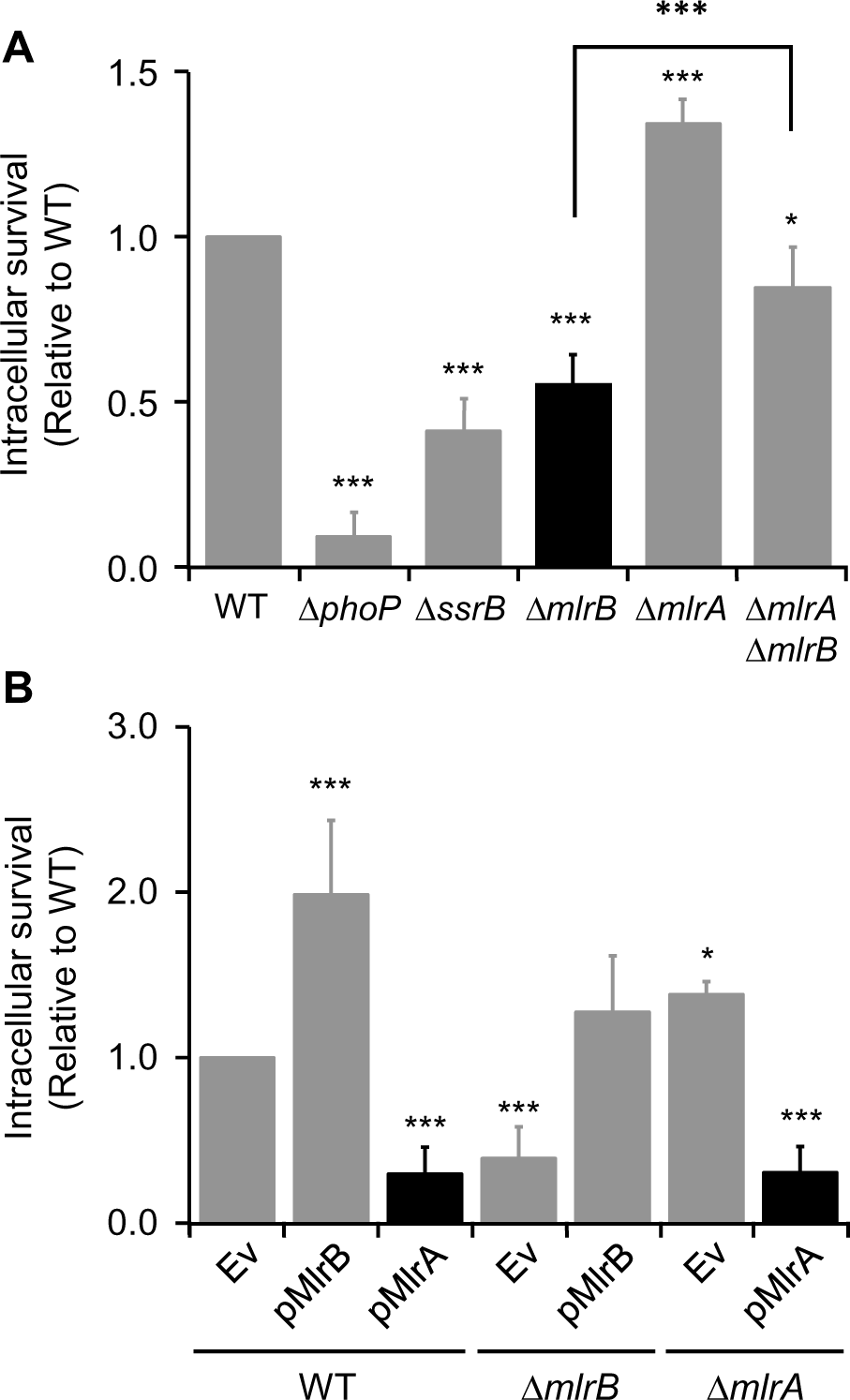
MlrB is required for intramacrophage survival. **A.** Survival of wild-type (WT), Δ*mlrB*, Δ*mlrA*, and Δ*mlrA* Δ*mlrB S.* Typhimurium strains in RAW 264.7 macrophages at 18 h after infection. The Δ*phoP* and Δ*ssrB* strains were included as controls. The values correspond to the average of at least three independent experiments carried out in duplicate and the error bars represent SD. **B.** Intramacrophage survival of wild-type (WT), Δ*mlrB*, and Δ*mlrA* strains ectopically expressing MlrB or MlrA from a multicopy plasmid was determined as in (A). (Ev) indicates the empty vector. The values correspond to the average of at least three independent experiments carried out in duplicate and the error bars represent SD. Asterisks denote statistical significance between means, with respect to WT. n.s., not significant; *, P < 0.05; **, P < 0.01; ***, P < 0.001.

*orf319* was not an intermediary in the MlrB-mediated intracellular survival, since its deletion did not alter the survival of either the wild-type or the Δ*mlrB* mutant (Fig. 5).

**Figure 5.**
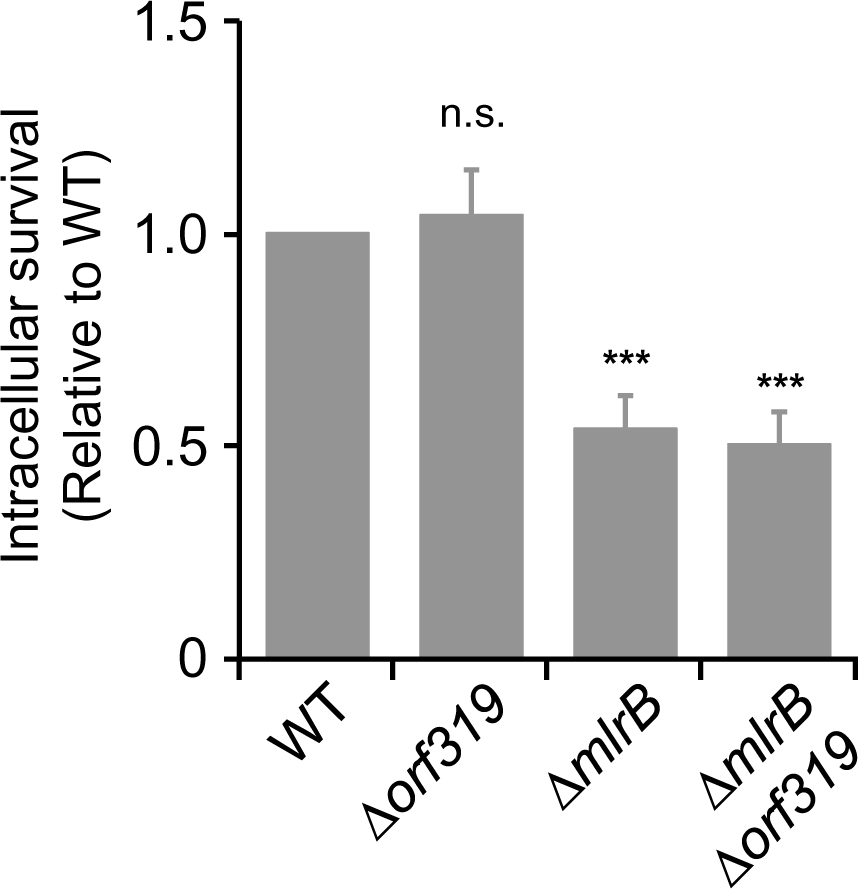
*orf319* inactivation does not affect intramacrophage survival. Survival of wild-type (WT), Δ*orf319*, Δ*mlrB*, and Δ*mlrB* Δ*orf319 S.* Typhimurium inside RAW264.7 macrophages at 18 h after infection. The values correspond to the average of at least three independent experiments carried out in duplicate and the error bars represent SD. Symbols above bars denote statistical significance between means, with respect to WT. n.s., not significant; *, P < 0.05; **, P < 0.01; ***, P < 0.001.

In view of the above results, and aware that MlrB did not affect *csgD* expression under laboratory conditions (Fig. 1), we hypothesized that MlrB could exert its function by modulating CsgD expression in a different context, i.e., within macrophages.

#### MlrB promotes intramacrophage survival by repressing *csgD* expression

We determined intramacrophage *csgD* transcription from a P*csgD*::*gfp* reporter at 0 (inoculum) and 18 h.p.i. A marked reduction of *csgD* transcription at 18 h.p.i. was observed in the WT strain but not in the Δ*mlrB*, which maintained similar values to that of the inoculum (Fig. 6A), indicating that, during *Salmonella* intramacrophage growth, MlrB is capable of negatively regulating *csgD* expression.

**Figure 6.**
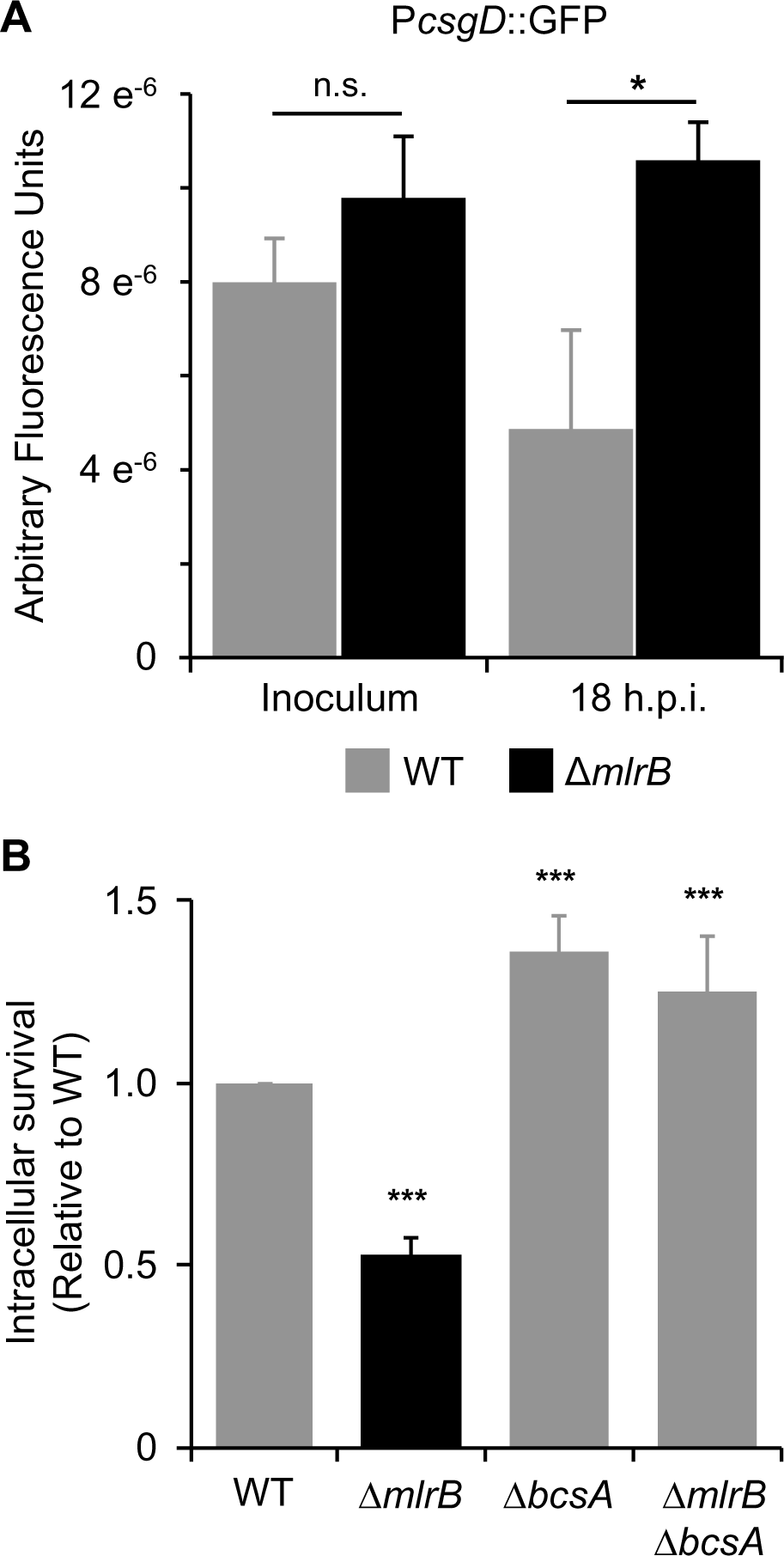
The defect in macrophage survival of the Δ*mlrB Salmonella* mutant depends on CsgD expression. **A.** CsgD expression increases inside macrophages in the absence of MlrB. Arbitrary fluorescence from a P*csgD*::*gfp* reporter plasmid expressed in wild-type (WT) or in *ΔmlrB* strains inside RAW264.7 macrophages at 0 (inoculum) and 18 h post infection. Values correspond to the ratio between fluorescence and the number of intracellular surviving bacteria. The data correspond to the average of at least three independent experiments carried out in duplicate and the error bars correspond to the SD. **B.** MlrB effect on *Salmonella* survival within macrophages is *bcsA* dependent. Relative survival of Δ*mlrB*, Δ*bcsA* and Δ*mlrB* Δ*bcsA strains* inside RAW264.7 macrophages at 18 h after infection compared to the wild-type (WT) strain. The values correspond to the average of at least three independent experiments carried out in duplicate and the error bars represent SD. In **A** and **B**, symbols above bars denote statistical significance between means, with respect to WT. n.s., not significant; *, P < 0.05; **, P < 0.01; ***, P < 0.001.

The repressive effect of MlrB on CsgD during intracellular survival suggests the involvement of MlrB in the control of EM production. This would explain the macrophage survival defect of the *mlrB* mutant. In particular, increased cellulose production, but not curli, has been reported to be detrimental for intracellular survival (Pontes *et al*., 2015). Indeed, deletion of the cellulose synthase coding gene, *bcsA*, restored wild-type intramacrophage replication levels of the Δ*mlrB* mutant (Fig. 6B).

Altogether these results indicate that MlrB, a SPI-2 encoded *Salmonella*-specific MlrA-like transcriptional regulator, links macrophage survival to extracellular matrix production.

## DISCUSSION

*Salmonella* biofilm formation is considered relevant for extra-host environments (Gerstel and Römling, 2003; Simm *et al*., 2014) or during gallbladder colonization in chronic infections (Crawford *et al*., 2010). Furthermore, although cellulose has been shown to be synthesized by *Salmonella* inside macrophages, its increased production is detrimental for survival within these cells and for systemic infection in mice (Pontes *et al*., 2015; Ahmad *et al*., 2016). Nevertheless, the biological significance of cellulose production by intra-macrophage bacteria remains to date poorly understood.

In this work we demonstrated that MlrB, a SPI-2 encoded MlrA-like transcription factor, is required for *Salmonella* intramacrophage survival. Further, our results indicate the MlrB negatively controls *csgD* expression and activation of the cellulose biosynthesis pathway when inside macrophages. We showed that, opposite to MlrA, MlrB expression increased under SPI-2 inducing condition (Fig. 2 and S2A). Consequently, we observed a 6-fold increase in MlrB levels in intramacrophages- grown *Salmonella* (Fig. 2C and Fig S4). Using *E. coli* MlrA target operators (Ogasawara *et al*., 2010) we identified another SPI-2 gene, *orf319*, as a MlrB- as well as MlrA-regulated gene (Fig. 3, S5 and S6), and showed that control of MlrB or MlrA over *orf319* depends on the growth conditions. For example, both MlrB and MlrA were active in repressing *orf319* transcription in LB (Fig. S6), but only MlrB repressed *orf319* under SPI-2 inducing conditions (Fig. 3).

On the other hand, and while only MlrA activated *csgD* expression in laboratory conditions (Fig. 1B), we showed an increased expression of *csgD* inside macrophages in the absence of MlrB (Fig. 6) indicating that in this environment MlrB acts as a repressor of the biofilm master regulator. In this sense, we propose a counteracting role between MlrB and MlrA inside macrophages. This is further supported by the increased survival of the Δ*mlrA* Δ*mlrB* double mutant strain compared with the mutant deleted in *mlrB* (Fig. 4A).

Our results sustain a MlrB-activating condition inside macrophages rather than a consequence of a balance between MlrB and MlrA concentrations because there was no MlrB-dependent effect on csgD expression either in LB or in InsPI-2 inducing conditions (Fig. 1B and Fig. S7). Nevertheless, this is under current investigation at the lab. Certainly, as the effector-binding C-terminal domains of MlrA and MlrB share low sequence identity (Fig. S1A), it is expected that each regulator would be modulated by different signals and therefore respond to different environmental cues.

The prediction of two MlrA-like operators in the *csgD* promoter may also account for the opposite role between MlrB and MlrA in controlling CsgD expression. Whether these regulators are interacting and recognizing the same sequence or they prefer one over the other needs to be investigated. In marked contrast, the only operator found in the promoter region of the MlrB- and MlrA-repressed gene *orf319* overlaps this gene’s -10 and -35 elements (Fig. S5B). It is feasible that binding of either MlrB or MlrA would hinder the RNA-polymerase from its productive interaction with the promoter. If this is the case, the proposed mechanism of action of these regulators differs from other *Salmonella* MerR-regulators like GolS or CueR (Checa *et al*., 2007; Pérez Audero *et al*., 2010; Humbert *et al*., 2013). Not only these canonical regulators do not impede the binding of RNA-polymerase, but they favor it (Pezza *et al*., 2016).

The deficiency in macrophage survival of the *mlrB* mutant strain can be attributed to an increased cellulose production, as the deletion of the cellulose synthase gene *bcsA* restored wild-type survival of the Δ*mlrB* strain (Fig. 6B) while the absence of Orf319, the other MlrB-controlled gene product, did not affect *Salmonella* survival inside macrophages (Fig. 5). This results is in concordance with the observation that overproduction of cellulose is the cause of the virulence deficiency of a Δ*mgtC* strain as the inactivation of *bcsA* restored wild-type virulence of the *mgtC* mutant (Pontes *et al*., 2015). MgtC is a virulence protein located in the inner membrane that controls ATP synthesis membrane potential by interacting with the α subunit of the F_1_F_0_ ATP synthase (Pontes *et al*., 2015), and phosphate uptake by inducing the PhoB/PhoR regulatory system (Lee *et al*., 2014). MgtC controls *bcsA* expression and also c-di-GMP intracellular concentration, which is an allosteric cellulose synthase activator, although the mechanisms involved in this regulation are still not known (Pontes *et al*., 2015).

Why cellulose production affects intramacrophage growth it is not known, although one possible explanation could be related to the glucose consumption that its production requires, that otherwise could be redirected to intramacrophage growth (Petersen *et al*., 2019). Cellulose production during *Salmonella* intramacrophage survival was associated to a persister subpopulation (Petersen *et al*., 2019). It would be interesting to know whether this slow replicating population is maintained as a safeguard reservoir and produces the EM component as a means of self-protection. In this sense, we propose that MlrB would act as a rheostat to balance intracellular growth versus presisters’ reservoir.

It has been shown that *Salmonella* serovars involved in systemic infection suffered an elevated genome degradation compared with the restricted gastrointestinal serovars although genome degradation is observed in all serovars analyzed (Nuccio and Bäumler, 2014). This correlates with the requirement of particular traits in each case, showing an increased loss of genes encoding Type III secreted effectors, fimbrial adhesins, and motility and chemotaxis genes in extraintestinal serovars compared to gastrointestinal pathovars (Nuccio and Bäumler, 2014).

Despite the lack of information about the gene-repertoire modulated during persistence either in the gallbladder or in *Salmonella*-containing granulomas, genes encoding biofilm formation components and its regulators appear to be conserved in both groups (Nuccio and Bäumler, 2014), suggesting that this trait is relevant for common processes.

In sum, our work describes the function of MlrB, a novel SPI-2-encoded, MlrA-like transcription factor required for *Salmonella* intramacrophage survival. Our genetic and biochemical studies allowed us to propose a plausible mechanism for this regulation, in which MlrB modulates extracellular matrix production when inside macrophages to regulate pathogenicity. Divergent patterns of expression and activities suggest that MlrB, which showed no obvious effect on regulating biofilm-formation *in vitro*, antagonizes MlrA *in vivo*. This not only illustrates how two closely related transcription factors counteract in response to specific environmental signals, but also contributes to the increasingly accepted notion that biofilm-formation is a complex process that affects both intra- and extra-host lifestyles.

## ACKNOWLEDGMENTS

We thank Gonzalo Tulin, Diego Serra and Susana Checa for critical reading and comments on the manuscript. pP*sseA*::*lacZ* plasmid was kindly provided by B. Bret Finlay. We are also grateful to María Dolores Campos, Marina Avecilla and Marina Perozzi for their excellent technical assistance. This work was supported by grants from Agencia Nacional de Promoción Científica y Tecnológica (grant PICT-2015- 2056) and the Spanish Ministry of Science and Innovation (BIO2016-77639-P).M.L.E. and N.R.F. are fellows of CONICET and L.V.H. is recipient of a Doctoral fellowship from ANPCyT. F.C.S. is a career investigator of CONICET and of the Rosario National University Research Council.

## AUTHOR CONTRIBUTIONS

MLE, NRF and FCS designed the experiments and wrote the manuscript. MLE, NRF, LVH and MGP performed the experiments. MLE, NRF, LVH, MGP, FGdP and FCS analyzed the data.

## MATERIALS & METHODS

### Bacterial strains and growth conditions

*S. enterica* serovar Typhimurium strains and plasmids used in this study are listed in Table S1. Oligonucleotides are listed in Table S2. Cells were routinely grown at 37°C in Luria–Bertani (LB broth) or on LB-agar plates, except when indicated. Ampicillin, tetracycline, kanamycin, and chloramphenicol were used when necessary at 100, 15, 50, and 20 μg ml^-1^, respectively. Culture media that emulate the intravacuolar environment used were PCN, InSPI-2 (Kröger *et al*., 2013) and LPM (Coombes *et al*., 2004).

All reagents and chemicals were from Sigma, except the Luria-Bertani culture medium that was from Difco. Oligonucleotides and enzymes were purchased from Life Technologies.

### Genetic and molecular biology techniques

The strains carrying gene deletions, chromosomal *lacZ* reporter fusions or 3xFLAG tags were generated by Lambda Red-mediated recombination followed by P22- mediated transduction using previously described protocols (Pérez Audero *et al*., 2010; Ibáñez *et al*., 2013; Pezza *et al*., 2016; López *et al*., 2018). When necessary, antibiotic resistance cassettes inserted at the deletion points were removed using FLP-mediated recombination (Datsenko and Wanner, 2000). DNA fragments as well as plasmids were introduced into bacterial cells by electroporation. All constructs were verified by DNA sequencing.

The Plasmid carrying the transcriptional fusion of the native *Salmonella* P_*csgD*_ promoter to *gfp* (Table S1) was constructed by cloning the PCR-amplified promoter into the SmaI site of pPROBE(NT) using previously described protocols (Miller *et al*., 2000).

### Quantification of biofilm adhesion to abiotic surfaces

To evaluate the adhesion to polystyrene microplates, a Crystal Violet (CV) binding and quantification protocol was implemented (Pitts *et al*., 2003) . For this, 150 μl of 1:100 dilutions of saturated cultures of the strains to be tested were deposited in 96- well polystyrene plates. The inoculated plates were incubated at the temperatures and times indicated. After incubation, the cultures were discarded and the wells were washed 4 times with distilled water by immersion and allowed to dry at room temperature. Then 200 μl of a 1% (w/v) aqueous solution of CV was added to each well and incubated for 20 minutes at room temperature. Subsequently, unbound CV was washed out by thorough immersion in distilled water. After drying the plates at room temperature, 200 μl of an ethanol:acetone mixture 80:20 (v/v) was added to each well. Desorption of the dye was allowed for one hour at room temperature on a shaking platform. Finally, the absorbance at 562 nm was recorded using a spectrophotometer (BioTek ELχ808).

### β-galactosidase activity assays

Measurements of β-galactosidase activity of strains carrying transcriptional fusions to the *lacZY* genes were made following a modification of the protocol proposed by Miller (Miller, 1972), and essentially as described in (Pérez Audero *et al*., 2010).

### Western blot analysis

Western blot analyses of 3xFLAG-tagged proteins or IgaA were carried out as described previously (Pontel and Soncini, 2009; Pérez Audero *et al*., 2010) with mouse anti-FLAG monoclonal (Sigma-Aldrich) or rabbit polyclonal anti-IgaA antibodies. Quantification of individual bands by densitometry was performed using the Image J Program, using the IgaA band as a load control (Lobato-Márquez *et al*., 2015).

### Eukaryotic cells culture conditions and gentamicin protection assay

*Salmonella* survival in RAW 264.7 macrophages was tested as described (Thompson *et al*., 2011). Briefly, macrophages were cultured in 24-well plates containing Dulbecco’s modified Eagle’s medium (DMEM) supplemented with 10% fetal calf serum (FCS) and were infected at a multiplicity of infection (MOI) of 10 bacteria per cell. The macrophage medium was supplemented with IPTG (1 mM) or Arabinose 0,05% (v/v) when these cells were infected with *Salmonella* strains harboring plasmid pUHE-21–2lacIq, pBAD30 or its derivatives. After infection, plates were incubated at 37°C for 30 min and then fresh D-MEM 10% FBS medium supplemented with gentamicin 100 µg/ml was added. After 1 h at 37°C, infected cells were incubated with medium containing gentamicin at a concentration of 30 µg/ml for a total of 18 h. At indicated time points, cells were washed and lysed with 0,1% Triton X-100 in PBS. Lysates were recovered and serially diluted. CFUs were determined on LB agar plates and the intracellular survival (fold of change) was determined.

For large-scale experiments needed to monitor protein production by intracellular bacteria, RAW 264.7 macrophages were incubated in 500 cm^2^ plates as described previously (Núñez-Hernández et al., 2013). Briefly, cells were seeded in BioDish-XL 500-cm^2^ plates until they reached confluence. Then they were infected with *Salmonella* at a MOI of 10 bacteria per cell. After 40 min, cells were washed tree times with PBS and then were incubated with fresh medium containing 100 µg/ml of gentamicin. After 1 h at 37 °C, infected cells were incubated with medium containing gentamicin at a concentration of 30 µg/ml until 18 h post infection. At indicated time points, infected macrophages were washed with PBS and lysed in a solution containing 4% SDS, 1% acidic phenol, and 19% ethanol in water. After 30 min of incubation at 4°C, intracellular bacteria were collected by centrifugation (27,500 g, 4°C, 30 min) and washed three times with 1 ml of a, 19% ethanol RNAse free solution. For western blot analyses, intracellular bacteria were processed as described previously (Pontel and Soncini, 2009; Pérez Audero *et al*., 2010). Immunodetection was carried out using mouse anti-FLAG monoclonal (Sigma-Aldrich) or rabbit polyclonal anti-IgaA antibodies (Lobato-Márquez *et al*., 2015).

### *csgD* intracellular expression

Measurement of *csgD* transcription from intracellular bacteria was performed using WT/pPromcsgD-gfp or ΔmlrB/pPromcsgD-gfp strains. A gentamicin protection assay was performed as described above, with modifications, as follows. RAW 264.7 macrophages were cultured in 6-well plates until they reached confluence. WT/pPromcsgD-gfp or ΔmlrB/pPromcsgD-gfp strains were grown ON at 37°C. At the indicated time point, invaded cells were washed four times with PBS and lysed with 0,1% PBS-Triton X-100. Lysates were collected, centrifuged 5 min at 6500 rpm and resuspended in PBS. Two hundred microliters of each sample were used to measure GFP fluorescence in a Synergy 2 Microplate Reader (Biotek) spectrophotometer (λ_exc_=485 nm, λ_em_=528 nm). One hundred microliters of each sample were used to determine the number of intracellular viable bacteria (CFU/ml) by serial dilution and plating on LB. Transcriptional induction values were expressed as fluorescence units per CFU.

### In silico analyses

A positional matrix of weights was generated through the MEME program (Bailey *et al*., 2009) to screen for MlrA targets in *S.* Typhimurium genome, using the reported *E. coli* MlrA-binding sequences in the promoters of *csgD, rplU-ispB, yrbA, dppB* and *cadC* (Ogasawara *et al*., 2010). This consensus was later confronted with the genome of *S.* Typhimurium LT2 using the FIMO tool (Grant *et al*., 2011).

### Statistical analysis

To test for statistical differences between means, one-way analysis of variance (ANOVA) and the Tukey multiple comparison test with an overall significance level of 0.05 were used. Calculations were performed with GraphPad Prism statistical software.

---

**Table S1.**
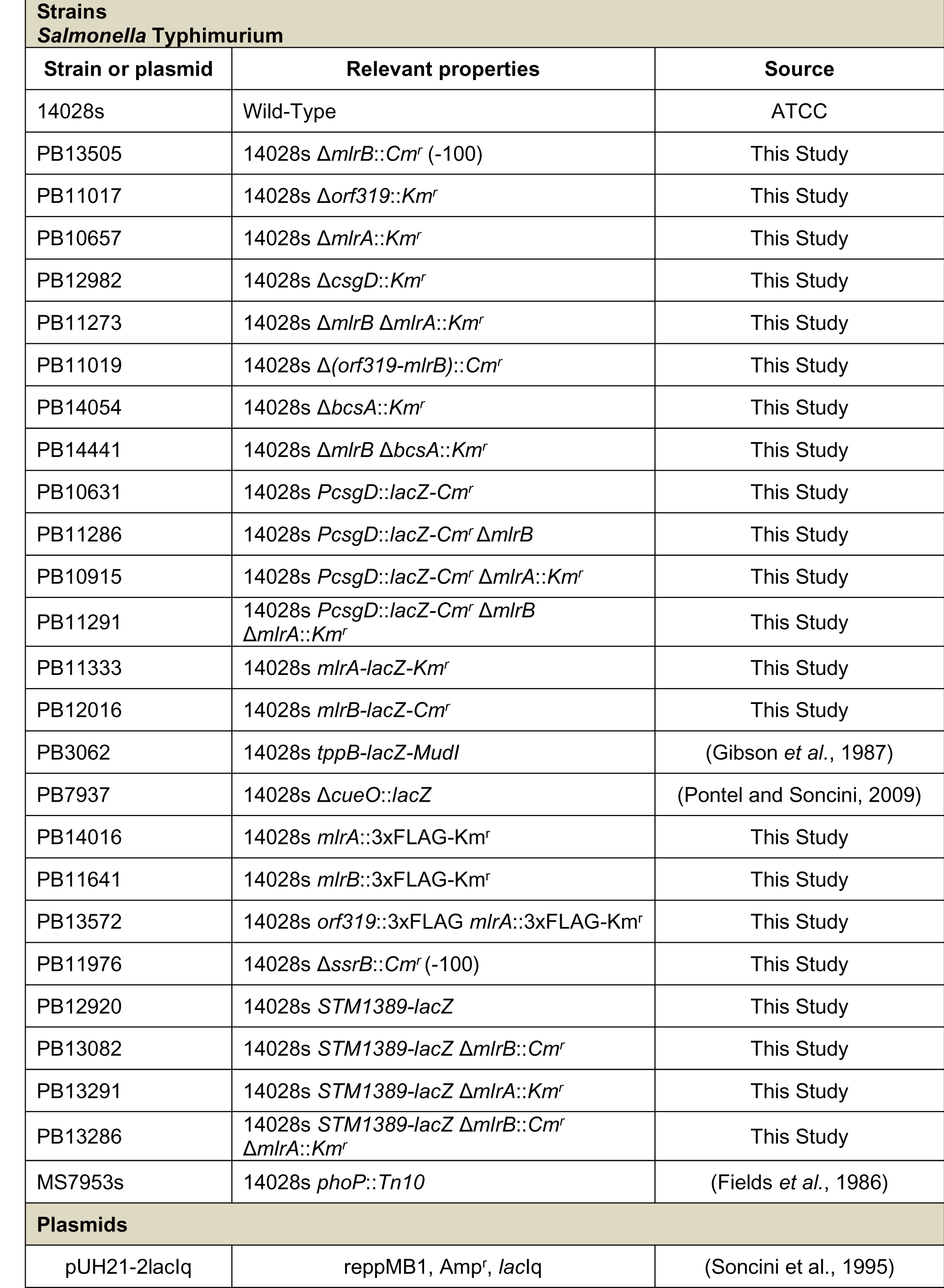
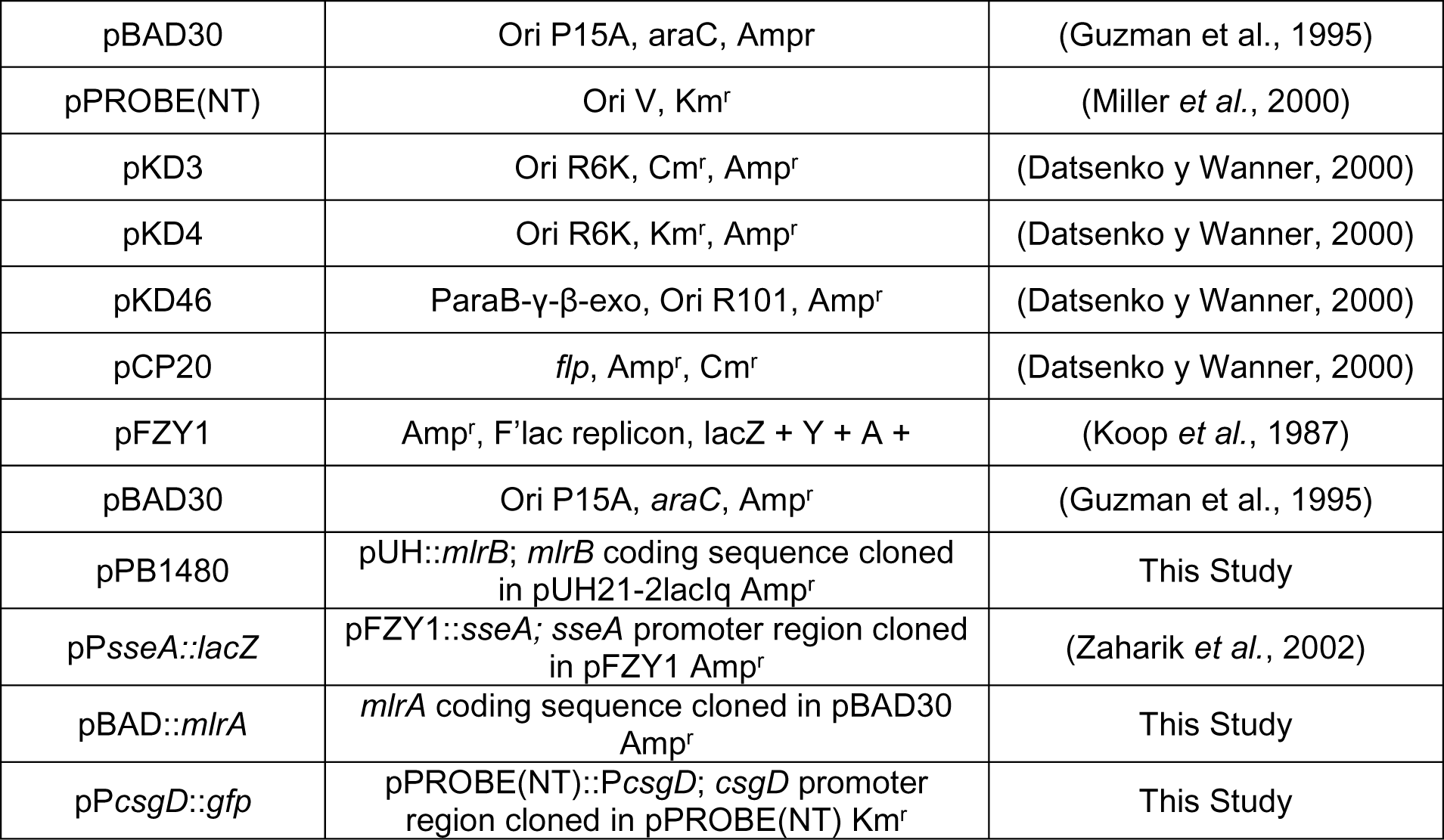
Bacterial strains and plasmids used in this study.

| **Strains** |  |  |
| --- | --- | --- |
| ***Salmonella* Typhimurium** |  |  |
| **Strain or plasmid** | **Relevant properties** | **Source** |
| 14028s | Wild-Type | ATCC |
| PB13505 | 14028s $\Delta mlrB::Cm^r$ (-100) | This Study |
| PB11017 | 14028s $\Delta orf319::Km^r$ | This Study |
| PB10657 | 14028s $\Delta mlrA::Km^r$ | This Study |
| PB12982 | 14028s $\Delta csgD::Km^r$ | This Study |
| PB11273 | 14028s $\Delta mlrB \Delta mlrA::Km^r$ | This Study |
| PB11019 | 14028s $\Delta (orf319-mlrB)::Cm^r$ | This Study |
| PB14054 | 14028s $\Delta bcsA::Km^r$ | This Study |
| PB14441 | 14028s $\Delta mlrB \Delta bcsA::Km^r$ | This Study |
| PB10631 | 14028s $PcsgD::lacZ-Cm^r$ | This Study |
| PB11286 | 14028s $PcsgD::lacZ-Cm^r \Delta mlrB$ | This Study |
| PB10915 | 14028s $PcsgD::lacZ-Cm^r \Delta mlrA::Km^r$ | This Study |
| PB11291 | 14028s $PcsgD::lacZ-Cm^r \Delta mlrB \Delta mlrA::Km^r$ | This Study |
| PB11333 | 14028s $mlrA-lacZ-Km^r$ | This Study |
| PB12016 | 14028s $mlrB-lacZ-Cm^r$ | This Study |
| PB3062 | 14028s $tppB-lacZ-MudI$ | (Gibson *et al.*, 1987) |
| PB7937 | 14028s $\Delta cueO::lacZ$ | (Pontel and Soncini, 2009) |
| PB14016 | 14028s $mlrA::3xFLAG-Km^r$ | This Study |
| PB11641 | 14028s $mlrB::3xFLAG-Km^r$ | This Study |
| PB13572 | 14028s $orf319::3xFLAG mlrA::3xFLAG-Km^r$ | This Study |
| PB11976 | 14028s $\Delta ssrB::Cm^r$ (-100) | This Study |
| PB12920 | 14028s $STM1389-lacZ$ | This Study |
| PB13082 | 14028s $STM1389-lacZ \Delta mlrB::Cm^r$ | This Study |
| PB13291 | 14028s $STM1389-lacZ \Delta mlrA::Km^r$ | This Study |
| PB13286 | 14028s $STM1389-lacZ \Delta mlrB::Cm^r \Delta mlrA::Km^r$ | This Study |
| MS7953s | 14028s $phoP::Tn10$ | (Fields *et al.*, 1986) |
| **Plasmids** |  |  |
| pUH21-2lacIq | reppMB1, Amp<sup>r</sup>, lacIq | (Soncini *et al.*, 1995) |
| pBAD30 | Ori P15A, araC, Amp<sup>r</sup> | (Guzman et al., 1995) |
| pPROBE(NT) | Ori V, Km<sup>r</sup> | (Miller *et al.*, 2000) |
| pKD3 | Ori R6K, Cm<sup>r</sup>, Amp<sup>r</sup> | (Datsenko y Wanner, 2000) |
| pKD4 | Ori R6K, Km<sup>r</sup>, Amp<sup>r</sup> | (Datsenko y Wanner, 2000) |
| pKD46 | ParaB-γ-β-exo, Ori R101, Amp<sup>r</sup> | (Datsenko y Wanner, 2000) |
| pCP20 | *flp*, Amp<sup>r</sup>, Cm<sup>r</sup> | (Datsenko y Wanner, 2000) |
| pFZY1 | Amp<sup>r</sup>, F'lac replicon, lacZ + Y + A + | (Koop *et al.*, 1987) |
| pBAD30 | Ori P15A, araC, Amp<sup>r</sup> | (Guzman et al., 1995) |
| pPB1480 | pUH::*mlrB*; *mlrB* coding sequence cloned in pUH21-2lacIq Amp<sup>r</sup> | This Study |
| pP*sseA*::*lacZ* | pFZY1::*sseA*; *sseA* promoter region cloned in pFZY1 Amp<sup>r</sup> | (Zaharik *et al.*, 2002) |
| pBAD::*mlrA* | *mlrA* coding sequence cloned in pBAD30 Amp<sup>r</sup> | This Study |
| pP*csgD*::*gfp* | pPROBE(NT)::P*csgD*; *csgD* promoter region cloned in pPROBE(NT) Km<sup>r</sup> | This Study |

**Table S2.**
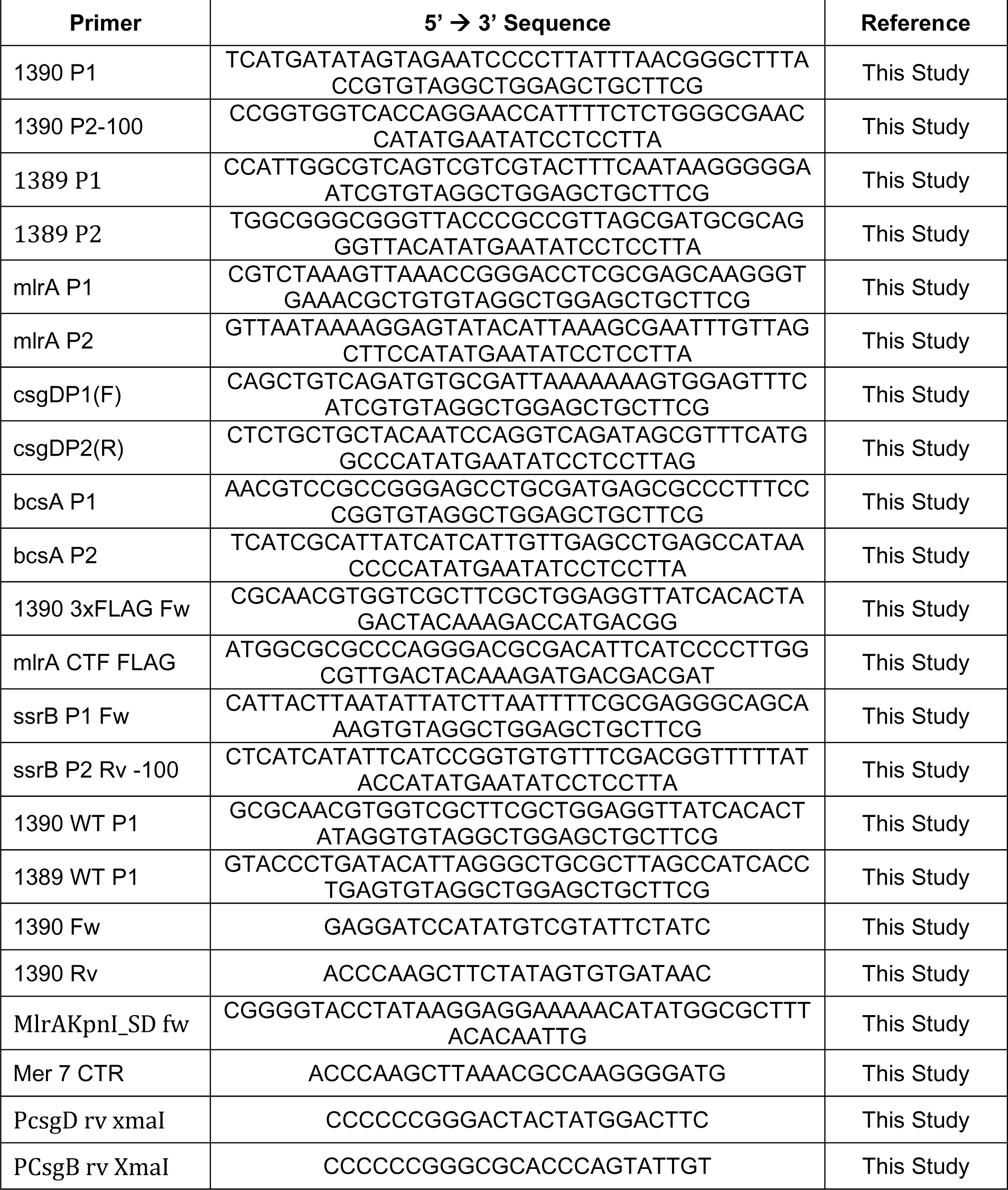
Oligonucleotides used in this study.

**Figure S1.**
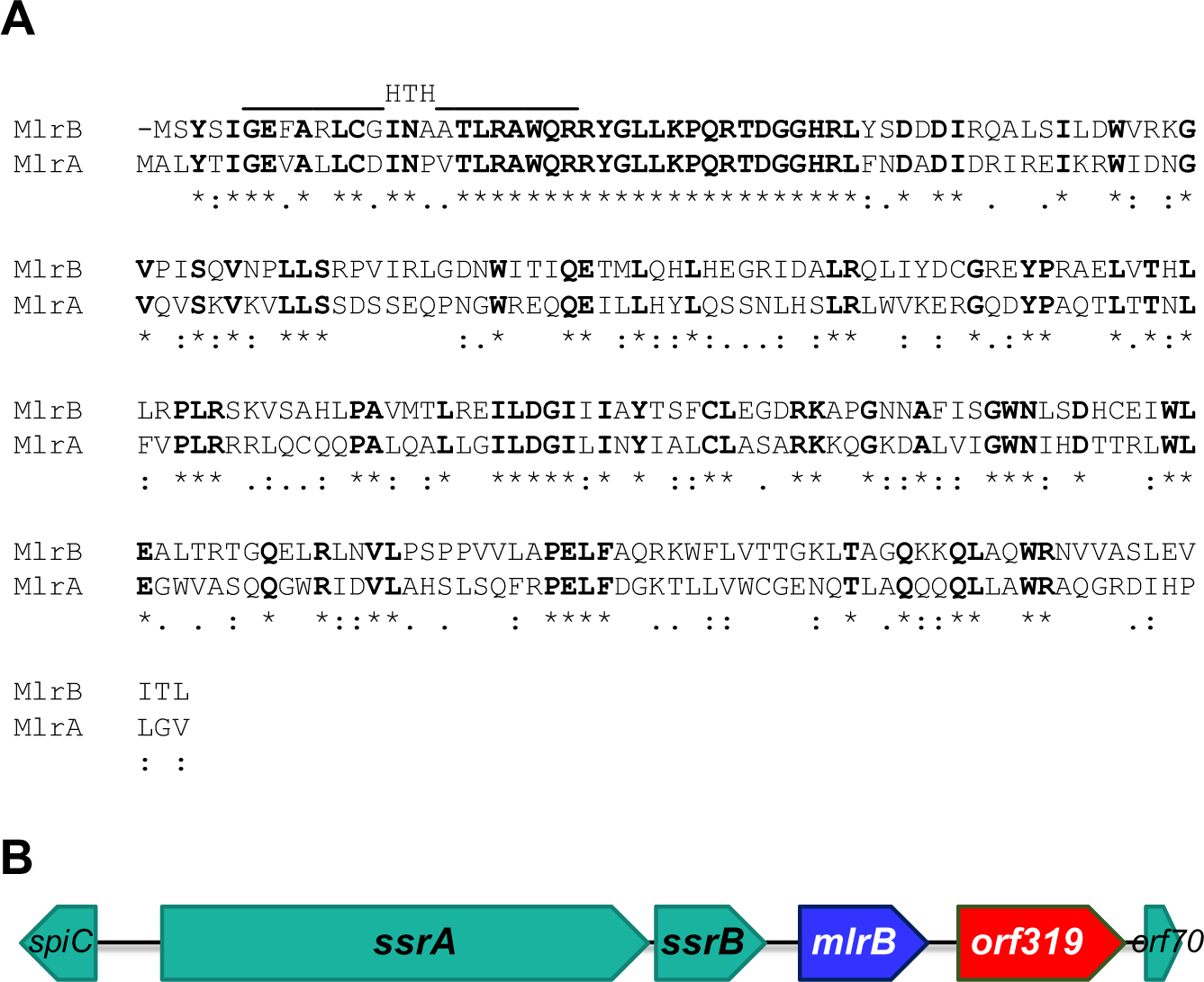
Amino acid alignment of MlrA and MlrB and *mlrB* genomic context. **A.** Alignment of amino acid sequences of MlrA (access ACY89107.1) and MlrB (access ACY88162.1) from *S.* Typhimurium strain LT2 (McClelland et al., 2001). Asterisks (*) indicate identical residues while dots (.) and double dots (:) indicate similarity. The predicted DNA-interacting helix-turn-helix region (HTH) is indicated (Humbert *et al*., 2013). **B.** Scheme of the SPI2 region containing mlrB and its neighbor genes.

**Figure S2.**
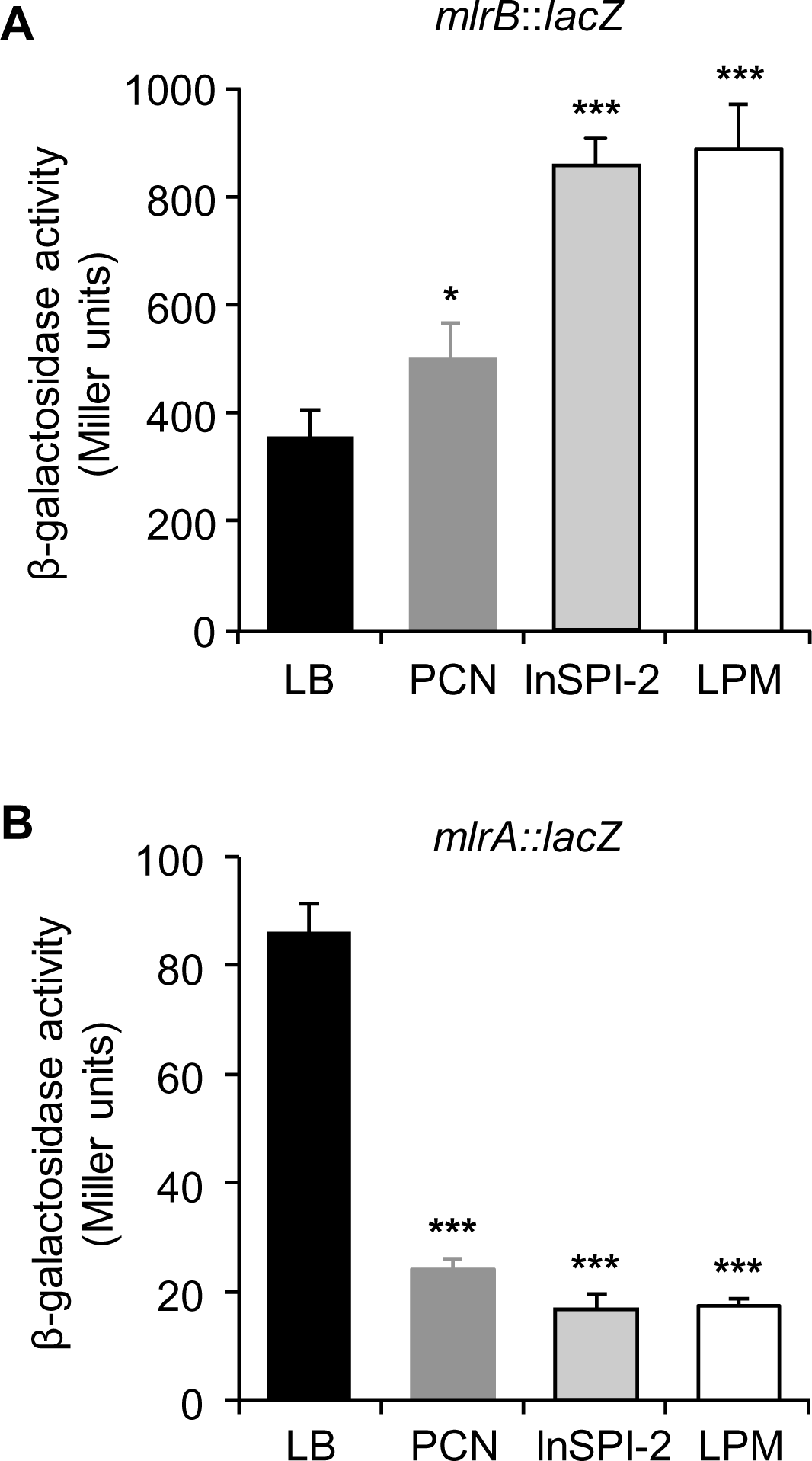
Comparative *mlrB* and *mlrA* expression in different growth media. β-galactosidase activity from **A**) *mlrB*::*lacZ*; or **B**) *mlrA*::*lacZ* transcriptional fusions, determined form cells grown to stationary phase in the indicated culture media, at 37°C. The values shown are the average of three independent experiments carried out in duplicate. Error bars correspond to the SD. Symbols above bars denote statistical significance between means, with respect to LB. n.s., not significant; *, P < 0.05; **, P < 0.01; ***, P < 0.001.

**Figure S3.**
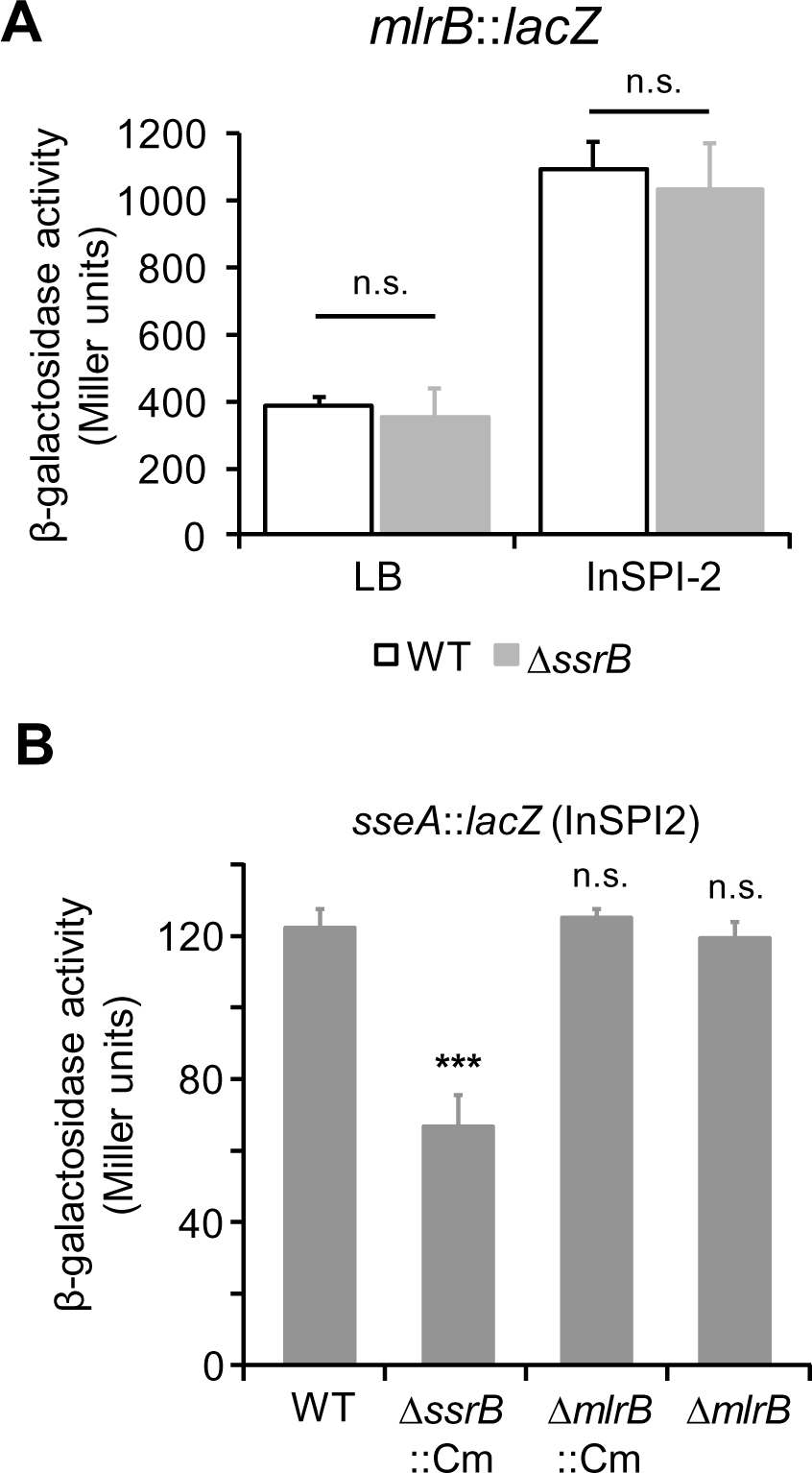
*mlrB* transcriptional induction under SPI-2 condition is not SsrA/SsrB-dependent. **A.** β-galactosidase activity of a *mlrB*::*lacZ* transcriptional fusion from wild-type (WT) or *ΔssrB*::*Cm* mutant cells grown overnight either in LB or in InSPI-2 culture media at 37 °C. Values correspond to the average of three independent experiments carried out in duplicate. Error bars correspond to the SD. **B.** β-galactosidase activity of a *sseA*::*lacZ* transcriptional fusion from wild-type (WT), *ΔssrB*::Cm (polar effect)*, ΔmlrB* or *ΔmlrB*::Cm (polar effect) strains. Cells were grown overnight in InSPI-2 media at 37 °C. The values represent the average of three independent experiments carried out in duplicate. Error bars correspond to the SD. In both cases, symbols above bars denote statistical significance between means as compared to the WT strain. n.s., not significant; *, P < 0.05; **, P < 0.01; ***, P < 0.001.

**Figure S4.**
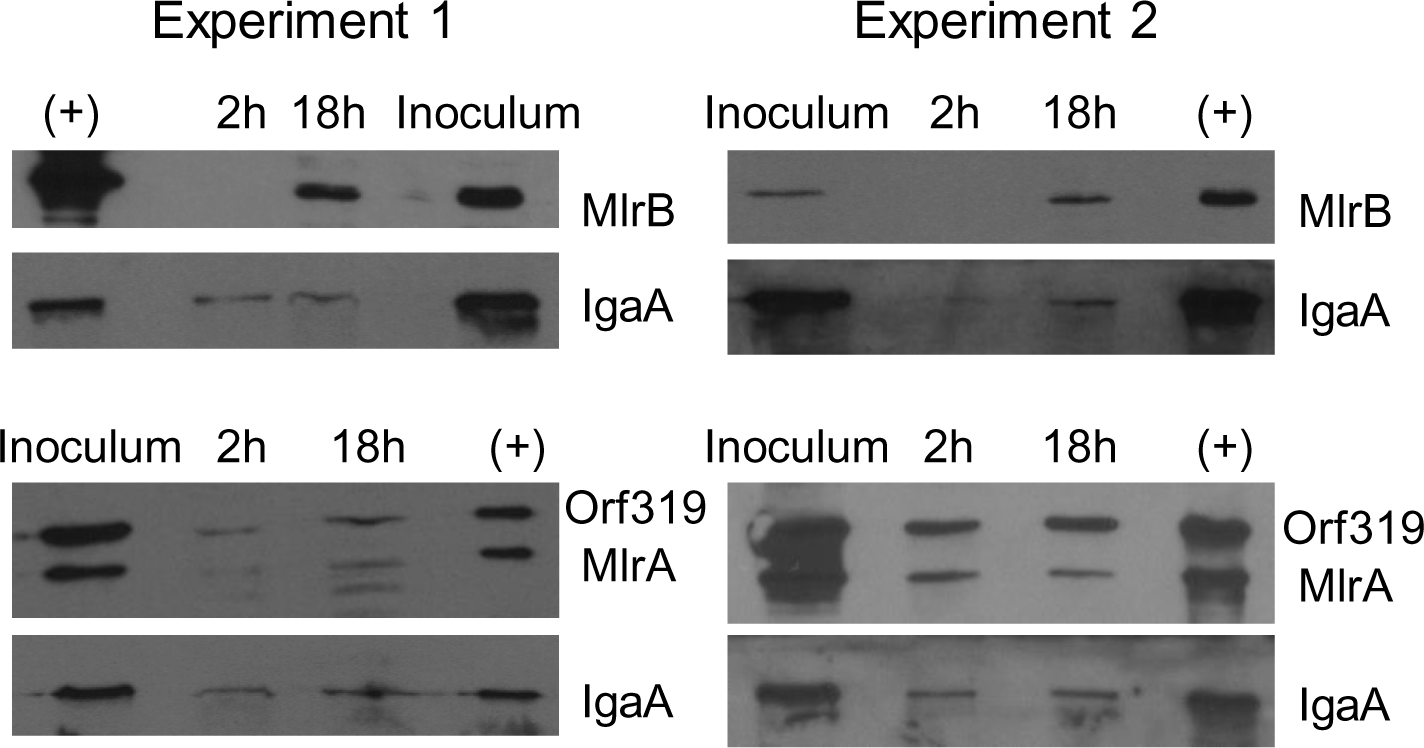
MlrB and MlrA protein levels inside macrophages. MlrB and MlrA protein levels detected by Western blots in total protein extracts obtained from *S.* Typhimurium *mlrB*::3×FLAG or *mlrA*::3×FLAG *orf319*::3×FLAG tagged strains grown inside RAW264.7 macrophages. The low-migrated bands in the MlrA Western blots correspond to Orf319-3xFLAG. The figure shows two different experiments. Anti- IgaA was used as load control.

**Figure S5.**
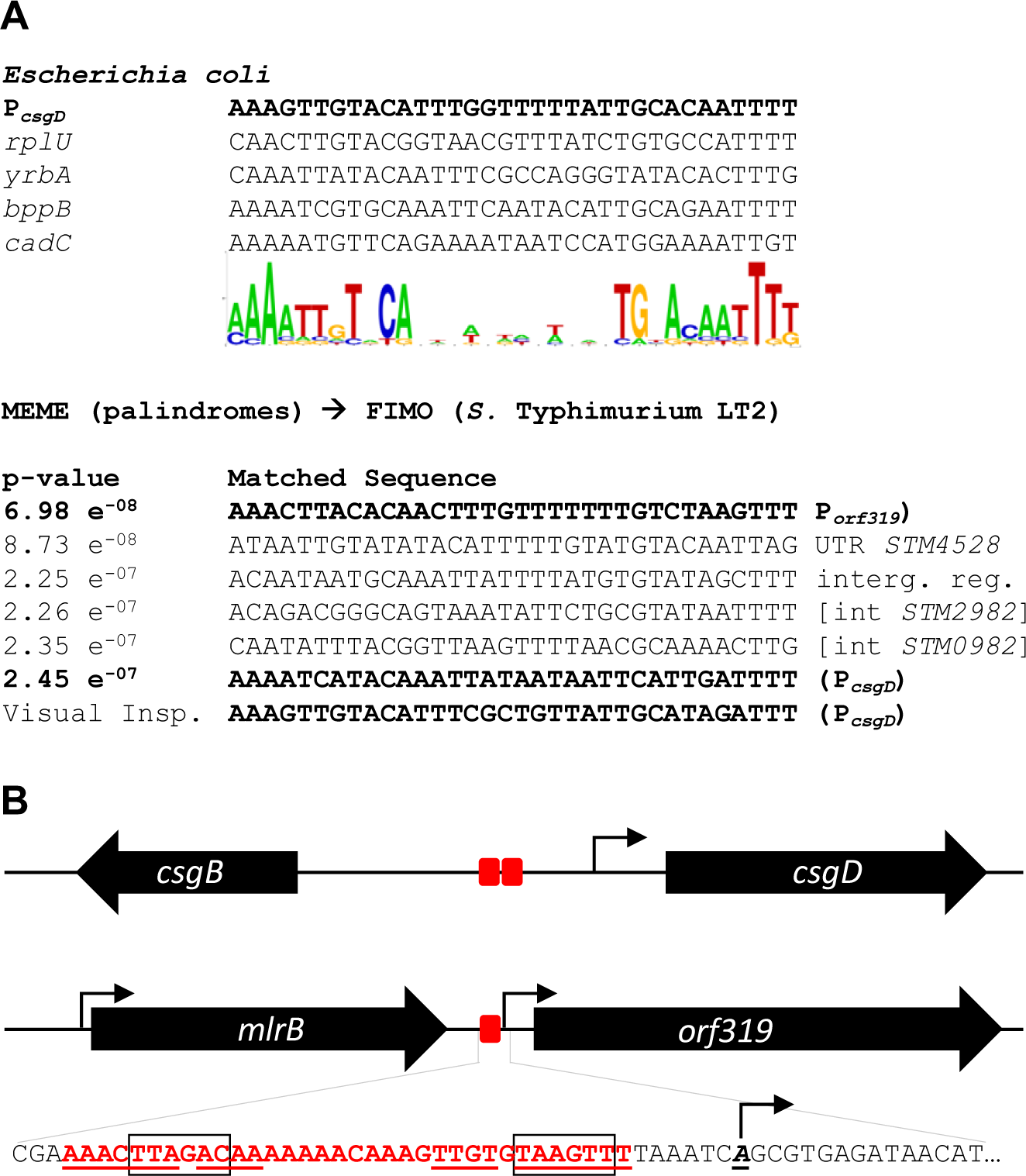
*In silico* screening for MlrA/MlrB DNA binding sites in the *S.* Typhimurium genome. **A.** A positional weight matrix was constructed using MlrA binding sequences reported in *E. coli* (Ogasawara *et al*., 2010) based on MEME/MAST, and used to screen the *S.* Typhimurium LT2 genome. Among the predicted MlrA-binding sites, the program detected one in the *csgD-csgB* intergenic region, and we identified a second by visual inspection. The program also identified a possible MlrA-binding site in the promoter region of *orf319*. **B.** Predicted MlrA/MlrB binding sites (red boxes) and *csgD* and *orf319* transcription initiation sites (arrows) (Kröger *et al*., 2013) are depicted. The DNA sequence of *orf319* promoter region is included. Boxes indicate the predicted -10 and -35 promoter elements. Underlined is the putative MlrB/MlrA recognized dyad.

**Figure S6.**
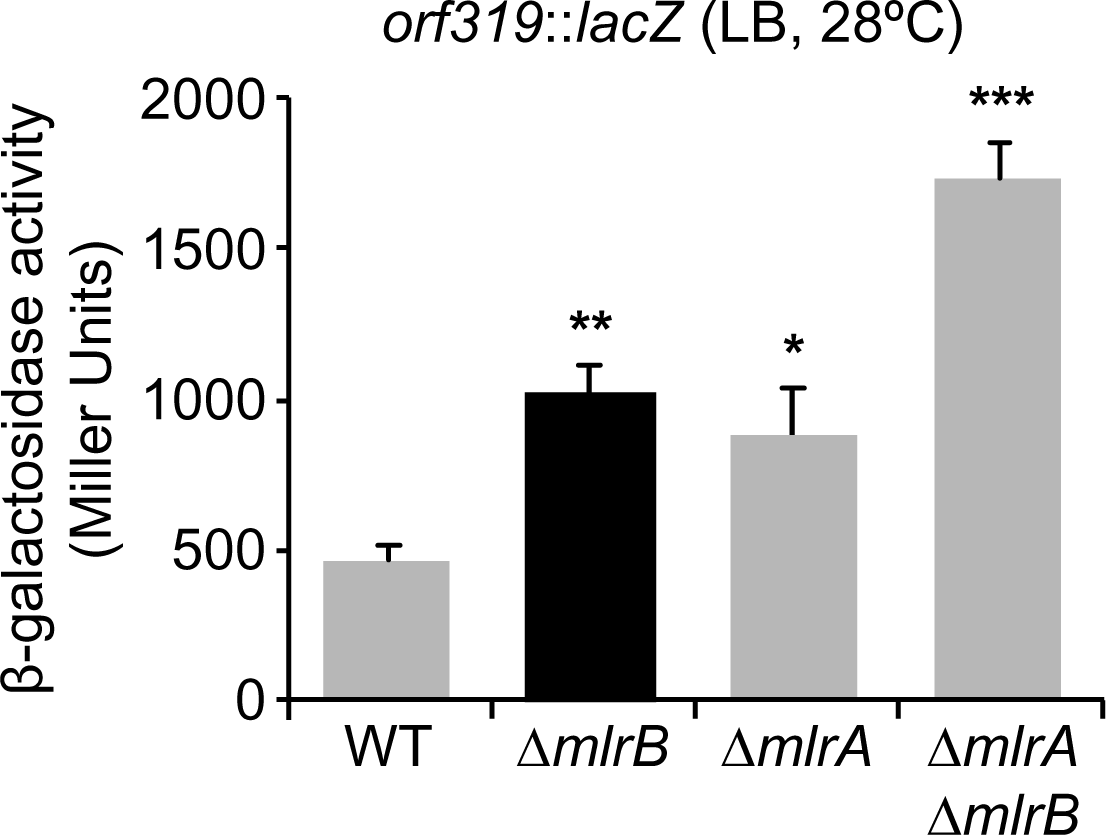
MlrB and MlrA co-dependent control of *orf319* transcription in LB. β-galactosidase activity of *orf319*::*lacZ* transcriptional fusion determined in wild-type (WT), Δ*mlrB*, Δ*mlrA*, and Δ*mlrA* Δ*mlrB S.* Typhimurium strains grown overnight in LB at 28°C. The values correspond to the average of three independent experiments carried out in duplicate and the error bars correspond to the standard deviation. Asterisks denote statistical significance between means, with respect to WT. *, P < 0.05; **, P < 0.01; ***, P < 0.001.

**Figure S7.**
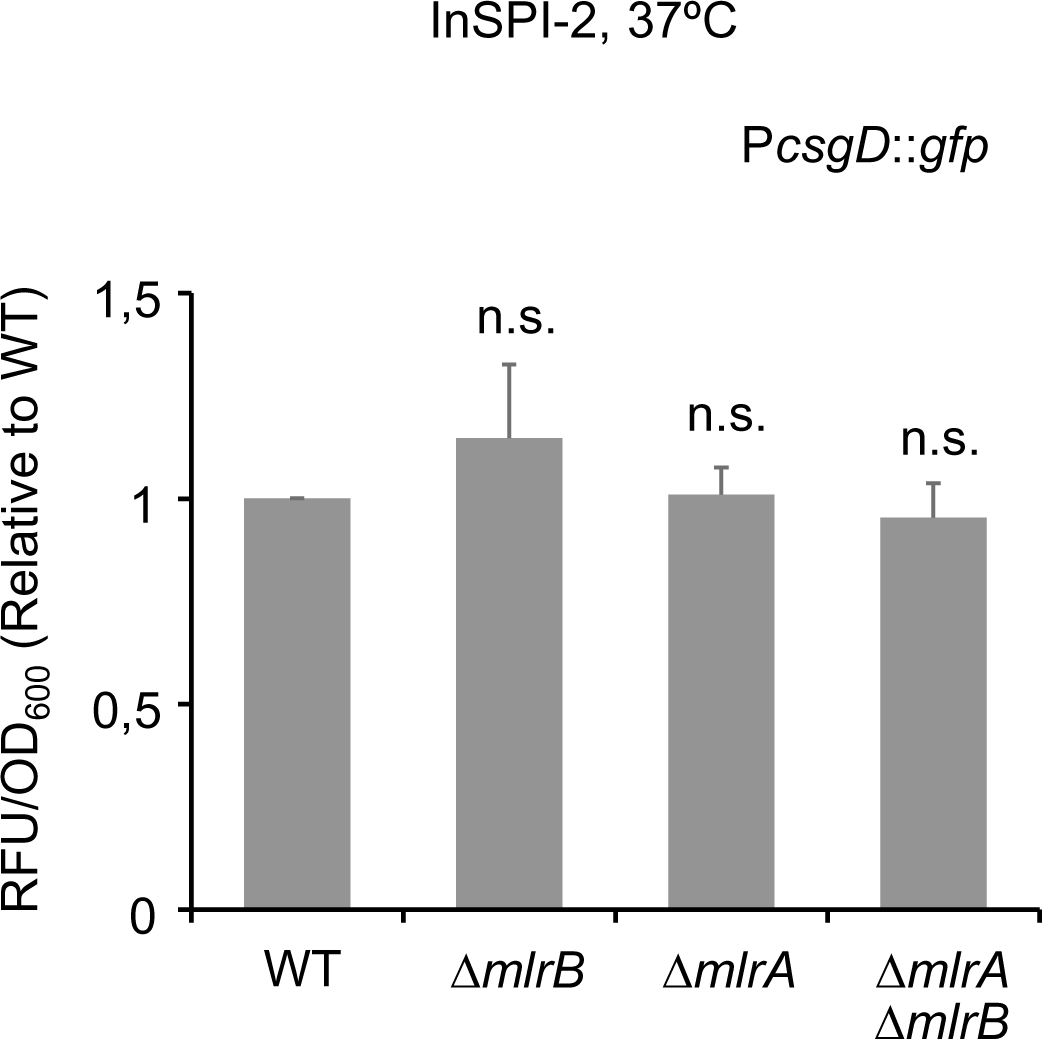
MlrB is not affecting CsgD expression under InSPI2 inducing conditions. Arbitrary fluorescence from a P*csgD*::*gfp* reporter plasmid expressed from wild-type (WT), *ΔmlrB*, *ΔmlrA*, or the *ΔmlrA ΔmlrB* strains grown at 37°C in InSPI-2. The data correspond to the average of at least three independent experiments carried out in duplicate and the error bars correspond to the SD. n.s., not significant.

